# Tracking wakefulness as it fades: micro-measures of Alertness

**DOI:** 10.1101/219527

**Authors:** Sridhar R. Jagannathan, Alejandro E. Nassar, Barbara Jachs, Olga V. Pustovaya, Corinne A. Bareham, Tristan A. Bekinschtein

**Author notes:** Present Address: Department of Psychology, University of Cambridge, Downing Street, Cambridge CB2 3EB, United Kingdom.

## Abstract

A major problem in psychology and physiology experiments is drowsiness: around a third of participants show decreased wakefulness despite being instructed to stay alert. In some non-visual experiments participants keep their eyes closed throughout the task, thus promoting the occurrence of such periods of varying alertness. These wakefulness changes contribute to systematic noise in data and measures of interest. To account for this omnipresent problem in data acquisition we defined criteria and code to allow researchers to detect and control for varying alertness in electroencephalography (EEG) experiments. We first revise a visual-scoring method developed for detection and characterization of the sleep-onset process, and adapt the same for detection of alertness levels. Furthermore, we show the major issues preventing the practical use of this method, and overcome these issues by developing an automated method based on frequency and sleep graphoelements, which is capable of detecting micro variations in alertness. The validity of the automated method was verified by training and testing the algorithm using a dataset where participants are known to fall asleep. In addition, we tested generalizability by independent validation on another dataset. The methods developed constitute a unique tool to assess micro variations in levels of alertness and control trial-by-trial retrospectively or prospectively in every experiment performed with EEG in cognitive neuroscience.

## 1. Introduction

Electroencephalography (EEG) has played a pivotal role in the non-invasive study of brain function (Niedermeyer and Silva, 2004). Typically in an EEG experiment the electrophysiological activity of the brain is recorded from the scalp of the participant while they are performing a cognitive task or under task-free conditions (e.g. resting state). In some task-based experiments, typically in the auditory or tactile domain, the participant performs the task with eyes-closed settings. Previous studies have shown that such eyes closed settings can create periods of momentary lapses of alertness (Barry et al., 2007). These periods are usually attributed to variable and long inter-trial intervals. The prevalence of this problem can be attested by studies mining large databases, which show that about a third of participants momentarily fall asleep in resting state conditions (Tagliazucchi and Laufs, 2014). Further, task-free settings such as mind wandering or simple non-active instructions can also lead to drowsiness and sleep (Goupil and Bekinschtein, 2012).

The above mentioned variations in alertness can usually be detected using variability in reaction times (Ogilvie, 2001). However in most of the EEG experiments such lapses are ignored and data confounded with drowsiness (or low alertness) are used for studying brain functions like attention and cognition. However, attention and many other cognitive sub-processes are known to be directly modulated by lack of alertness in normal (Bareham et al., 2014; Chennu and Bekinschtein, 2012) as well as clinical populations (Dobler et al., 2005). Hence, fluctuations in alertness need to be measured, to include or exclude trials of low/high alertness to adequately test predefined hypotheses. This argument is illustrated with an experiment in Figure 1.

**Fig 1:**
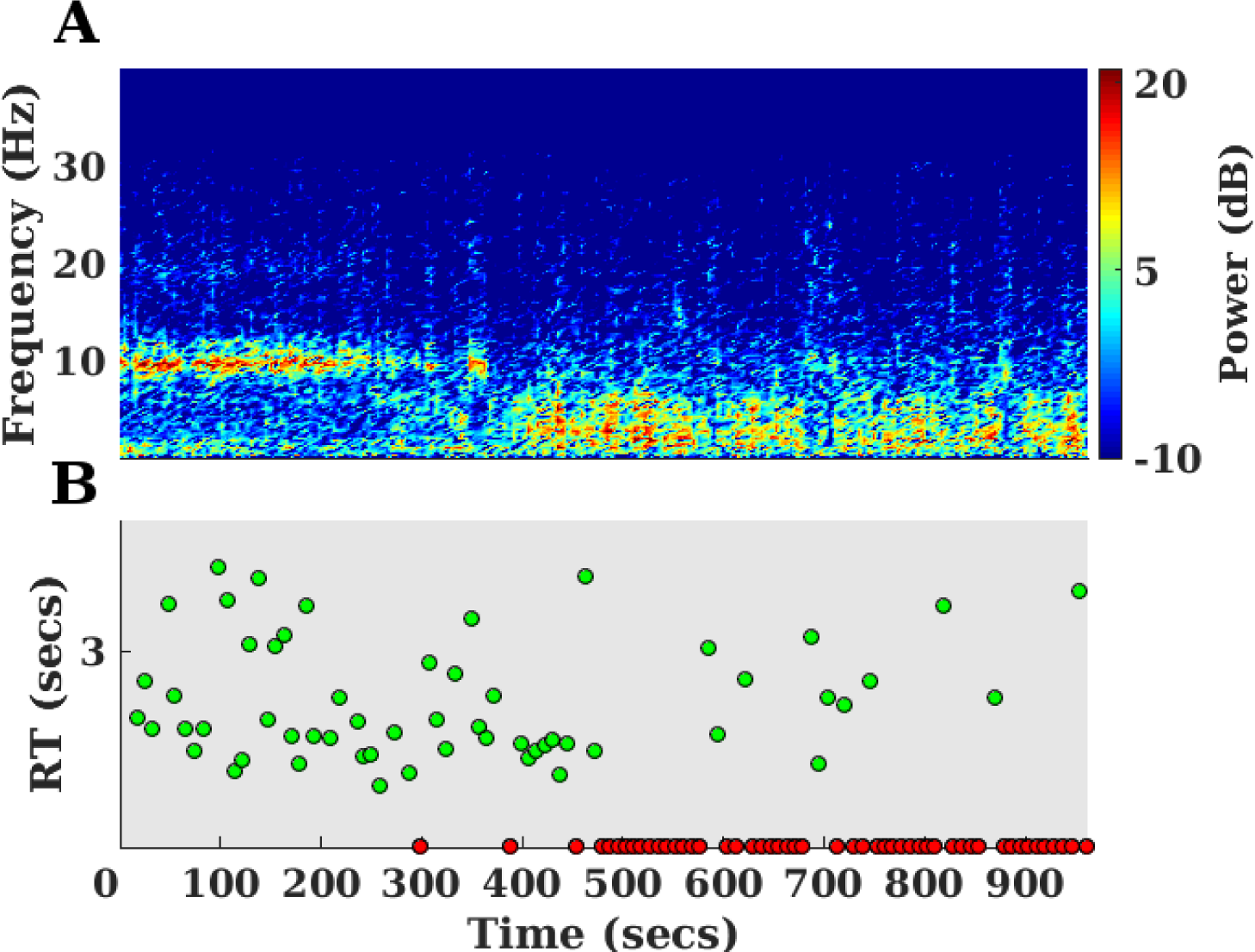
Differing alertness levels indicated by frequency profile changes and reaction time variability during an auditory experiment in a sample participant. (A) Depicts the changes in the power level in different frequency bands in the Occipital electrodes in the pre-trial period of an auditory experiment at different time points. (B) Reaction times at trials presented along the different time points in the same experiment. The variability in the reaction times (B) and thus reduction in alertness levels closely follows the change in the frequency profile (A) from alpha (8-12 Hz) to theta (6-8 Hz)

Figure 1(B) shows a typical EEG experiment (Kouider et al., 2014) where the participant responds to auditory stimuli while having their eyes closed. In the beginning of the experiment the participant responds to the stimuli in a reliable manner (green dots) by less variation in reaction times. As time progresses the reaction times become more variable and the participant intermittently fails to respond (red dots). This variation is also captured in the frequency profile of the EEG (occipital sites) during the pre-trial periods of the task as depicted in Figure 1(A). When the participant responds reliably, the frequency profile shows power in the alpha range (8-12 Hz) and as they become drowsy the alpha power disappears and low frequency power in the theta range (6-8 Hz) increases. Thus the frequency profile preceding the trial could predict the variability in the responses. In other words, such spectral changes can be used to detect the momentary lapses in alertness that causes variability in the reaction times.

The typical techniques that are used to clean or remove the data from such drowsiness contaminated episodes would be to score the above mentioned pre-trial periods using traditional sleep scoring techniques (Berry et al., 2012). These scoring techniques depend on the frequency profiles described earlier. However, they face multiple problems. Firstly, sleep scoring techniques rely on having at least 30 sec of data (Berry et al., 2012). However in most cognitive experiments the pre-trial periods last at most 4-5 sec. Secondly, automated methods (Tagliazucchi et al., 2012) that are validated using such sleep scoring techniques classify data into American Association of Sleep medicine (AASM) based sleep stages like wakefulness, N1, N2 etc. But such momentary lapses of alertness require more fine grained scoring techniques that operate on a smaller time range with different features capable of capturing micro variations in alertness levels. Finally, some techniques use the simple variation in reaction times mentioned earlier to capture moments of low alertness. But this suffers from the problem of longer reaction times being confounded by other factors such as task difficulty (Bareham et al., 2014).

Thus the above mentioned problem of fluctuations in alertness requires a novel solution. Our proposal is to tackle the problem in the following manner: Firstly, we identify these alertness contaminated episodes, through the use of Hori scale (Tanaka et al., 1996) that captures the micro variations in alertness. Though the prime purpose of the Hori system is to identify and characterise the sleep onset process, it contains features that enable us to identify variations in levels of alertness in more fine grained durations (4 sec) compared to traditional sleep scoring using wakefulness, N1 and N2. Secondly, we use human scorers to identify different levels of alertness using the Hori scale on a dataset where the participants are allowed to fall asleep while performing the task. Thirdly, we show that despite the clarity of the Hori scale, it is impractical to perform, time consuming and difficult to learn, as elucidated by the low degree of agreement among human scorers. Fourthly, we produce a practical solution to this problem using an automated technique (involving Support Vector Machine - SVM and individual element detectors) and compute performance measures by training and testing the algorithm on a dataset labelled by gold standard ratings (converging Hori ratings from multiple scorers). Finally, to estimate the reliability and generalisability of our method, we tested the same in another independent dataset to show its utility.

This paper is organized as follows. In the first section, we describe the method of using the Hori scale using human scorers and provide an overview of the automated method. In the second section, we evaluate and scrutinise the results of the human scorers with agreement measures and motivate the use of automated algorithm using validation measures. In the final section, we discuss the developments made in this paper and produce concluding remarks on the usefulness of the method developed here.

## 2. Materials and methods

### 2.1. Participants and datasets

The first dataset (herein Dataset#1) consisted of 20 native English speakers performing a semantic categorization task while falling asleep (Kouider et al., 2014). The task consisted of listening to words that belonged to a particular semantic category (e.g. animals or objects) and classifying them accordingly using a left or right button press. Each trial consisted of an auditory stimulus (spoken word: animal or object) presented binaurally with an intertrial interval of 6-9 sec.

The second dataset (herein Dataset#2) consisted of 31 participants performing an auditory masking task while falling asleep (Noreika et al., 2017a). The task consisted of listening to a target sound (e.g. beep) that was randomly masked by different noise duration. Participants reported whether they heard the target using a button press. Each trial consisted of an auditory stimulus (target) sometimes masked by noise, presented binaurally. The next trial was presented after a pause of 8-12 sec after the response or 13-17 sec (in case of no response).

In both the experiments subjects were seated on a reclining chair in a dark room and were permitted to fall asleep during the task. The participants were also evaluated on the basis of Epworth Sleepiness scale (Johns, 1991) and only easy sleepers were recruited.

### 2.2. EEG acquisition

Dataset#1: EEG was recorded using 64 Ag/AgCl electrodes (NeuroScan labs) with Cz as reference. The electrode impedances were kept below the recommended levels of the manufacturer. The signal was acquired at a sampling rate of 500 Hz.

Dataset#2: EEG was recorded using 129 Ag/AgCl electrodes (Electrical Geodesics Inc) with Cz as reference. The electrode impedances were kept below 100 KΩ. The signal was acquired at a sampling rate of 500 Hz.

### 2.3. Pre-processing

EEG data was pre-processed with custom made scripts in MATLAB (MathWorks Inc. Natick, MA, USA) using EEGLAB toolbox (Delorme and Makeig, 2004). The data was filtered between 1 and 30 Hz and was then resampled to 250 Hz. Furthermore, it was epoched from 4000ms to 0ms to the onset of the stimuli. Bad channels were then detected if the activity in the spectrum of the channel exceeds ±4 standard deviation of overall activity in all channels. The detected bad channels were then interpolated using spherical interpolation, after which trials that exceed the amplitude threshold of ±250uV were removed in a semi automatic fashion. The amplitude threshold was liberal as K-complexes usually exceed ±150uV.

Before proceeding to use the above datasets for scoring using the Hori scale it would be pertinent for us to first introduce the Hori system of scoring and inform the readers about the augmentations made in the system to suit the current purpose of measuring changes in alertness levels.

### 2.4. Hori Scale

Hori and colleagues subdivided the sleep onset process into 9 different substages (Tanaka et al., 1996). The first two Hori stages (1,2) correspond to wakefulness. The next six Hori stages (3-8) correspond to the sleep stage N1. The last stage of Hori (9) corresponds to the beginning of N2 sleep (Iber et al., 2007).

Here we decided to augment classical Hori stages with another stage (10) that would correspond to the appearance of K-complexes. The rationale behind this addition is the appearance of K-complexes definitively mark the entrance to N2 sleep. While spindles can still serve this purpose, their variability in duration and disagreement among human raters (Warby et al., 2014) motivates the use of K-complex. The following is a brief description of the elements in the hori scale based on (Ogilvie, 2001) and are shown in Figure 2.

**Fig 2:**
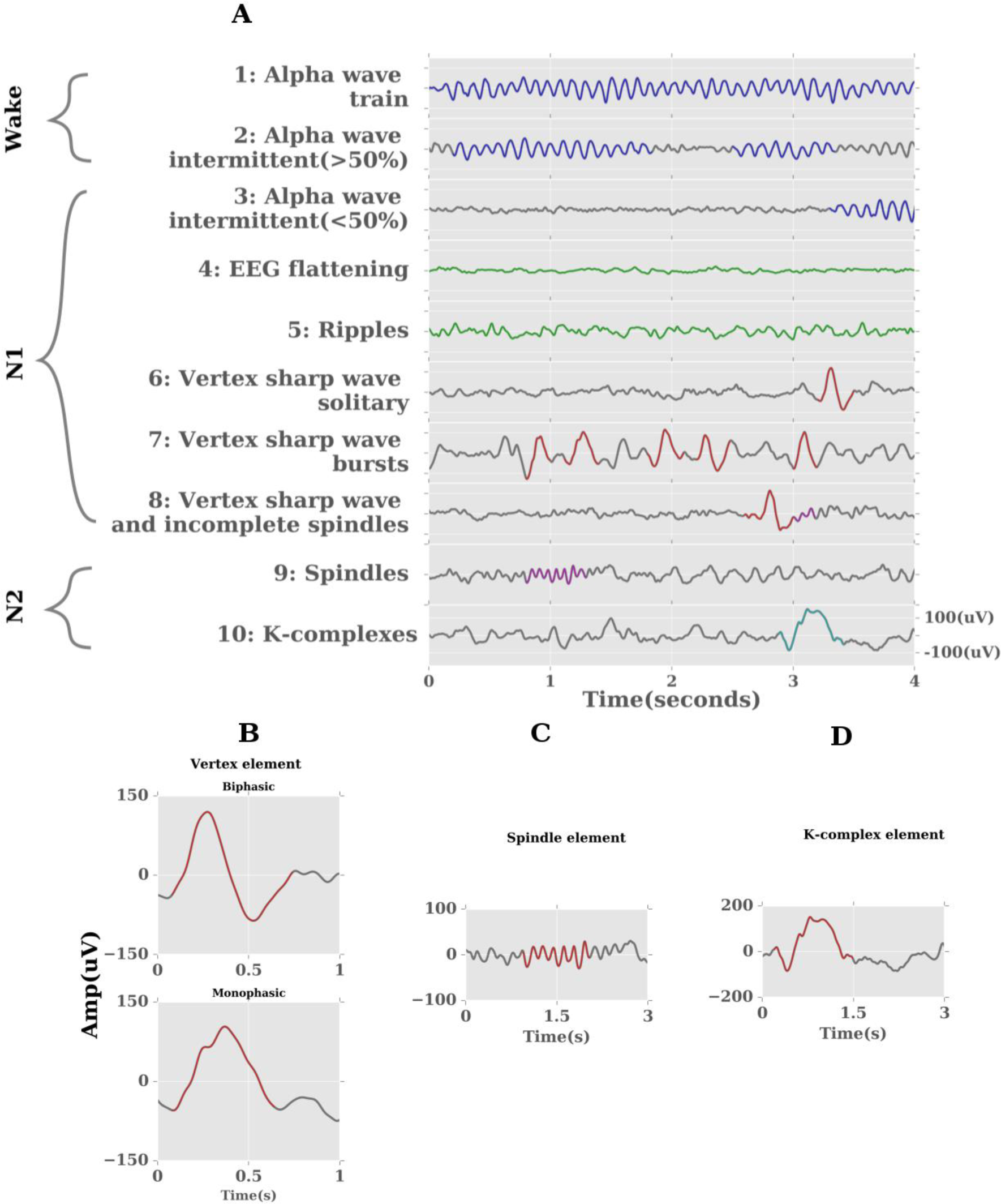
(A) Modified Hori scale for detecting differing alertness levels using EEG. The grey waves indicate background activity and coloured regions indicate characteristic elements for respective Hori stages. AASM based sleep stage classification is also represented for compatibility to classical sleep scoring. Grapho-elements of Hori scale in detail: (B) Vertex sharp waves: Biphasic consists of a sharp negative deflection followed by a positive one, whereas Monophasic consists of only a sharp negative deflection. (C) Spindles: transient patterns with frequency (12-16 Hz) and minimum duration of 0.5 sec. (D) K-complex elements: sharp positive deflection followed by a larger negative one with a duration of at least 0.5 sec

#### 2.4.1 Alert elements

##### Alpha waves

Alpha waves are elements that occur in the range of 8-12 Hz during relaxed wakefulness. They are more pronounced in the eyes closed condition, when the participant is transitioning from alert to relaxed wakefulness (Hori 1-2). Alpha elements are usually more pronounced in EEG from occipital regions.

*<u>Hori 1</u>*: Epoch is composed of only alpha wave trains (at least 20uV).

*<u>Hori 2</u>*: Alpha wave trains occupy more than 50% (but less than 100%) of the activity in the epoch.

#### 2.4.2 Drowsy elements

##### Alpha waves

Alpha activity usually decreases when the participant transitions from relaxed wakefulness to drowsy (Hori 3).

##### Theta waves

Theta waves are elements that occur in the range of 3-8 Hz. They have relatively higher amplitudes than the alpha elements and characterise the transition to N1. Theta activity is usually pronounced in the central and temporal regions (Hori 5).

*<u>Hori 3</u>*: Alpha wave trains occupy less than 50% of the activity in the epoch.

*<u>Hori 4</u>*: Activity flattening without any clear element (amplitude < 20 uV).

*<u>Hori 5</u>*: Low voltage theta waves (ripples) with amplitude between 20 uV-50 uV.

#### 2.4.3. Grapho elements

##### Vertex sharp waves

Vertex waves are grapho elements that occur in the beginning of the transition to sleep (Hori 6-8). Appearance of them indicates an altered state of responsiveness in the cerebral cortex (Rodenbeck et al., 2006). The vertex waves can be either monophasic or biphasic. In both cases there is usually a sharp negative discharge followed by a positive one. In the case of biphasic waves, the amplitude of the positive components should be at least 50% of the negative component and at most equal to the level of the negative component. The amplitude of the vertex sharp waves is found to be maximal in parietal and frontal regions (Cz based reference).

*<u>Hori 6</u>:* Epoch containing only one well defined vertex sharp wave.

*<u>Hori 7</u>*: Epoch containing more than one vertex sharp wave.

##### Spindles

Spindles are grapho elements that occur in the beginning of the transition to stage N2 of sleep (Hori 9). They are regarded as transient patterns of EEG activity with a frequency of 12-16 Hz with a minimum duration of 0.5 sec (complete spindles). Spindles in general should be distinguishable from the background activity. The typical waxing and waning of spindle shape is vital to distinguish the pattern from high alpha activity. The spindles were found to be prominent in temporal and frontal regions (Cz based reference).

*<u>Hori 8</u>*: Contains at least one vertex wave and an incomplete spindle (<0.5 sec).

*<u>Hori 9</u>*: Contains one well defined spindle (>0.5 sec).

##### K-complexes

K-complexes are grapho elements that occur in the N2 stage of sleep (modified Hori 10). It starts with a sharp positive wave followed by a large negative wave. The duration of the initial negative wave should be smaller than the positive wave. The overall duration of the K-complex must be at least 0.5 sec. The K-complexes were found to be prominent in frontal, temporal and parietal regions (Cz based reference).

*<u>Hori 10</u>:* Contains at least one well defined K-complex.

### 2.5. Manual Hori-scoring

For the purpose of manually scoring each epoch according to the Hori scale, the EEG data was further low pass filtered below 20 Hz and only 21 channels (Fig. 3(A)) derived using the standard 10-20 system were evaluated. The details of manual scoring is as follows:

**Fig 3:**
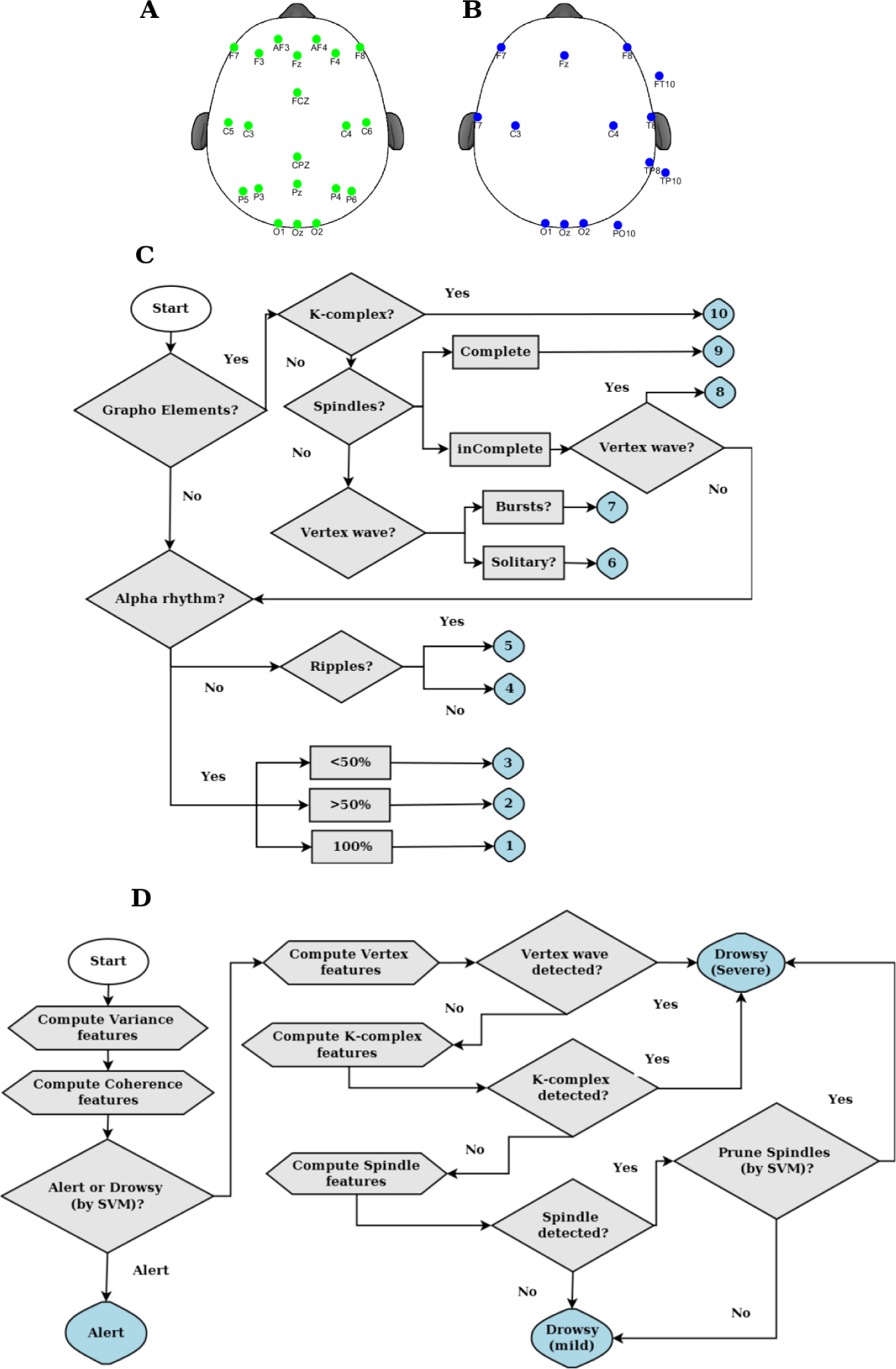
(A) Electrode sites used for manual Hori scoring based on 21 channels of the locations mainly derived from 10-20 electrode sites. (B) Electrodes used for automatic algorithmic method based on sampling from locations in Occipital, Central, Temporal, Parietal, Frontal regions. (C) Step by step technique to manually score each trial using the Hori scale. The preliminary step involves identifying presence of grapho-elements followed by specific identification of k-complexes, spindles and vertex waves. In the absence of grapho-elements, the trials are scored with identification of alpha rhythms. (D) Brief flow chart of the automatic algorithm. The preliminary step involves computation of the predictor variance and coherence features, followed by identification of alert and drowsy trials using SVM. Further, drowsy trials are identified into specific grapho-elements using detectors of elements like vertex, k-complex, spindles.

Dataset#1: Each pre trial epoch (-4000 to 0ms) was rated independently by 3 raters. Of which one was an experienced electrophysiologist (rater C) and 2 of the other raters (A, B) had learnt the technique immediately prior to scoring them independently. All participants were scored by the 3 raters, except for one participant that was scored only by raters A and B. As data from all participants was used based on consensus rule developed in section 2.6.1 this did not affect the results in anyway.

Dataset#2: Each pre trial epoch (-4000 to 0ms) was rated independently by 1 rater and was further verified with another experienced rater. One participant was ignored from further analysis as the original trial order could not be recovered from the raw EEG data.

The raters in dataset#1 scored each trial based on a manual algorithm depicted in Fig 3(C). The rater in dataset#2 scored each trial based on the description provided in (Ogilvie, 2001).

### 2.6. Automatic method

The automatic algorithm was first developed and tested using Dataset#1 and then independently validated using Dataset#2.

#### 2.6.1. Group consensus rule: creation of gold standard dataset

Before training and testing the algorithm, we decided to create labels in our input data (Dataset#1) that can be used by our algorithm for supervised learning. In our case, we decided to create a gold standard label for each trial that is based on a group consensus rule. For this purpose, we first subdivided the Hori ratings of each epoch per rater into Alert (Hori: 1,2), Drowsy-mild (Hori: 3,4,5), Drowsy-severe (Hori: 6,7,8,9,10). The gold standard label was computed using a simple majority among the raters. If there was no consensus, then the corresponding trials were ignored from further analysis. This group consensus rule was used in Dataset#1 and each trial was labelled into Alert, Drowsy (mild), Drowsy (severe). The creation of this gold standard dataset ensured that the algorithm was trained and tested with trials that were unambiguous and non-spurious.

#### 2.6.2. Electrode Choices

The electrodes depicted in Fig 3(B) were chosen for computing the various features used in different steps of the algorithm. The electrodes were chosen in such a way that we sample the Occipital, Frontal, Central, Parietal, Temporal regions. Furthermore, the choices were motivated for maximising the signal to noise ratio for the given reference electrode (Cz).

Dataset#1: Occipital: Oz, O1, O2; Frontal = F7, F8, Fz; Central = C3, C4;

Parietal = Pz; Temporal = T7, T8, TP8, FT10, TP10;

Dataset#2: Occipital: E75, E70, E83; Frontal = E27, E123, E11; Central = E35,

E110; Parietal = E90; Temporal = E109, E101, E115, E100;

A brief flow chart of the automatic algorithm is shown in Fig 3(D).

#### 2.6.3. Support Vector Machines

The first step in our algorithm involves computing features that are capable of distinguishing the various levels of alertness in the data. After which the features are used to devise a classifier capable of separating the Alert (Hori:1-2) from Drowsy (Hori: 3-10). We decided to use Support vector machines for this part of the classification as the classification problem is guaranteed to converge to an optimal solution (Platt, 1998; Tagliazucchi et al., 2012).

Support vector machines (SVM) are a class of supervised learning models. Formally, SVM consists of building a hyperplane or a set of hyperplanes in a high dimensional space with the criteria to maximise the distance of separation between the closest data (train-data) point of any class (functional margin) (Cortes and Vapnik, 1995). The choice of such a functional margin would lower the generalization error for new data points (test-data). The motivation to map the data onto higher dimensional space is driven by the fact that most often the classes are inseparable in the lower dimensional space (Boser et al., 1992). The mapping to higher dimensional space is achieved by the use of a kernel function *k*(*x, y*).

The kernel function avoids the need to compute individual data points in the transformed data space (computationally expensive) by using the euclidean inner product (kernel trick). In our paper, we used the MATLAB interface of the open source machine learning library (LIBSVM) (Chang and Lin, 2011) that supports use of kernel SVMs for nonlinear mappings. We used the

Radial Basis Function (RBF) as our kernel *k*(*x*,*y*) = e^(−γ‖x−y‖^2^^).

For training the classifier to produce optimal performance (accuracy) we need to select the optimal value of (γ, C). γ controls the curvature of the hyperplane and C represents the penalty parameter for the soft-margin. Parameter selection is achieved by performing a grid search in (γ, C) in the space 2^−1^,..,2^225^. We could not perform a leave one participant out cross validation, as this would produce an overfitting of parameters as different people fell asleep in different ways (proportion of alert, drowsy(mild), drowsy(severe) trials). Hence the data from all participants was collated and then divided into 5-folds (Tagliazucchi et al., 2012). Each of the 5-folds was made using stratified sampling such that the overall representation of sub-classes remained similar in each fold. This will avoid the problems of over-representation prevalent while using random-sampling. The first four folds were used to train the classifier to choose the parameters (γ, C) and the last fold was used to test the same. In order to measure the performance of the classifier we decided to use sensitivity, specificity, f1-score.

The definition of the performance measures used are as follows:

*<u>Accuracy:</u>* This is defined as the number of correctly classified data points divided by the overall number of classifications made.

*<u>Sensitivity:</u>* This refers to the ability of a classifier to correctly detect the true class among the classifications made. It is obtained by the (TP/TP+FN). It is also known as recall. TP: True Positives, FN: False Negatives.

*<u>Specificity:</u>* This refers to the ability of a classifier to correctly ignore the classes that don’t belong to the true condition. It is obtained by (TN/TN+FP). TN: True Negatives, FP: False Positives.

*<u>F1-score:</u>* This is the harmonic mean between precision and recall. Precision refers to measure of exactness of classifier. It is obtained by (TP/TP+FP). Recall refers to the sensitivity of the classifier.

As the input data contains different kinds of features, it was scaled using the minimum value and range before applying the SVM.

#### 2.6.4. Feature Computation

To use the above mentioned SVM for classification we need to compute the following features that can allow the classifier to distinguish between different classes.

##### Predictor Variance

The EEG data in the occipital region was first decomposed into time-frequency for each spatial sample (electrode) per epoch (-4000 to 0ms pre-trial). Predictors for each epoch were then generated based on the variations in the spectral power of the frequency bins A:[2-4 Hz], B:[8-10 Hz], C:[10-12 Hz], D:[2-6 Hz] per epoch. The predictors were then fit to the data per electrode-epoch and the variance explained is computed per electrode-epoch.

The first step is to transform the data *x*[*n*] into time-frequency representation (predictors) using the formula below, where *n* represents time domain with 1 ≤ *k* ≤ *N*

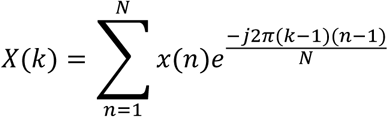

The next step is to compute the power in the transformed representation

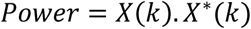

Followed by computing the predictor variance

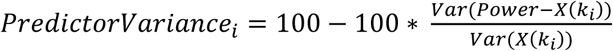

Where *i* represents the frequency band index (A,B,C,D) and *Var* represents the residual variance. Intuitively the predictor variance tries to capture the variance in the signal explained by different frequency bands and the SVM later on uses this feature for classification.

##### Coherence

Coherence was computed per trial in the electrodes in the occipital, frontal, central, temporal regions in the frequency bins: Delta:[1-4 Hz], Theta:[4-7 Hz], Alpha:[7-12 Hz], Sigma:[12-16 Hz], Gamma:[16-30 Hz]

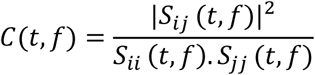

Where *C*(*t*, *f*) represents the coherence value at trial *t* and frequency band *f*

*S_ij_* represents cross power spectral density between signal *i* and *j*

*S_ij_*, *S_jj_* represents auto power spectral density.

After the detection of the drowsy trials using the above mentioned features, the following detectors are used to further subclassify them into drowsy (mild) and drowsy (severe).

#### 2.6.5. Grapho element detectors

##### 2.6.5.1. Vertex-wave-detectors

Both monophasic and biphasic waves were detected using the parietal electrodes. The signal was first resampled to 100 Hz and then filtered from 0.25-6 Hz. After which the signal in each trial was further scaled with respect to its minima. Peaks that are above a specific threshold are then detected and the negative peaks are used to classify the elements as mono or biphasic (algorithmic, parametric details described in supplementary methods)

##### 2.6.5.2. Spindle detectors

Spindles were detected using the temporal electrodes. The signal was first resampled to 100 Hz and then a continuous wavelet transform using morlet function as the mother wavelet was applied. The coefficients of this transform are then normalized and then further provided a rank according to the magnitude. Each rank is further normalized to compute the probability of the spindle occurrence at each time point. Further spindle locations are pruned using a snapshot of the detected location (algorithmic, parametric details described in supplementary material).

##### 2.6.5.3. K-complex detectors

K-complexes were detected using all the electrode sites in Fig 3(B). The signal was first resampled to 100 Hz and then filtered from 0.25-6 Hz. After which the signal in each trial was further scaled with respect to its maxima. Peaks that are separated by at least 1.5 sec and below a specific threshold are then detected. Further to which peaks above a specific threshold in the next 1.5 sec are detected. The positive peak should be at least half of the magnitude of the negative (algorithmic, parametric details described in supplementary material).

In summary a total of 32 features (12 from predictor variance; 20 from coherence) are used in the first stage detection of alert trials from drowsy trials. After the drowsy trials are parsed by the element detectors, the spindle elements are pruned again by a separate SVM using the same 32 features as above (depicted in Figure 3(D)).

## 3. Results

### 3.1. Manual Hori-scoring

In order to measure the reliability of scores given by the 3 different raters on different subjects in Dataset#1 we used two different measures of inter-rater agreement (Fig 4).

**Fig 4:**
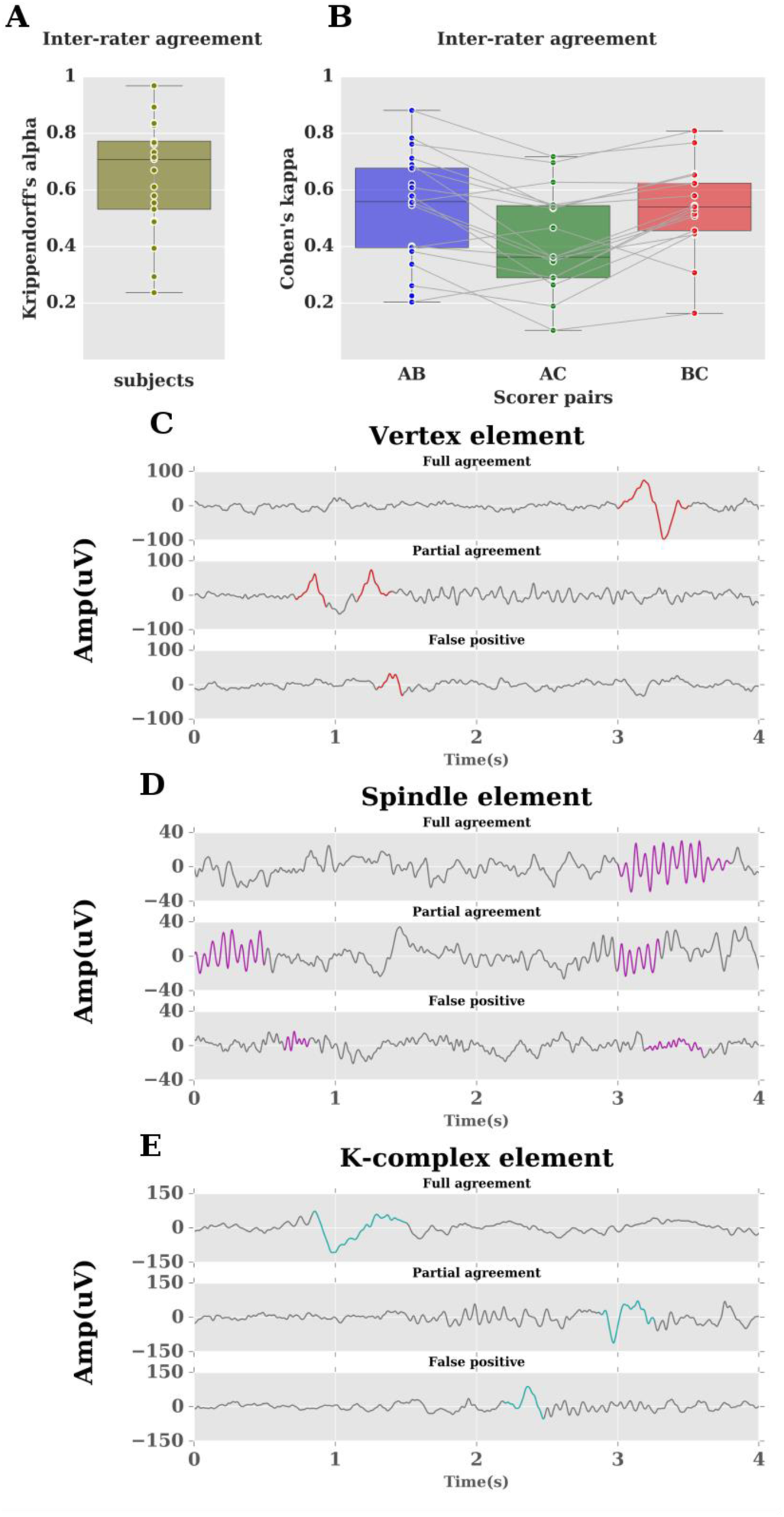
Inter-rater agreement among different scorers (A,B,C). (A) depicts agreement measured using Krippendorffs alpha. Each data point refers to score from a single subject. (B) depicts agreement measured using Cohen’s kappa. Each data point refers to kappa scores from a single subject based on a pair of two different scorers. Inter-rater disagreement is typically caused due to misclassification of Grapho elements: (C) depicts typical Vertex wave agreement/disagreement among scorers highlighted in red. (D) depicts typical Spindle element agreement/disagreement among scorers highlighted in magenta. (E) depicts typical K-complex agreement/disagreement among scorers highlighted in cyan. Full agreement refers to cases where all 3 raters agree, Partial agreement refers to cases where 2 of them agree, and false positives refer to cases where at least one of the rater misclassifies an element.

Firstly, we used Krippendorff’s alpha to compute the agreement between the 3 raters (A, B, C) per subject of Dataset#1. In general alpha scores of above 0.8 are reliable and those between 0.8 and 0.667 can only be used to draw tentative conclusions (Giannantonio, 2010). We can observe from Fig 4(A) that at least 9 subjects are below 0.667 (mean 0.65) indicating the unreliable nature of scoring each subject among raters. Secondly, we used Cohen’s kappa score (weighted) to measure the degree of inter-rater agreement between pairs of raters (AB, AC, BC) of Dataset#1. In general kappa values of above 0.8 are considered strong, between 0.8 and 0.4 as strong to weak, below 0.4 as poor (McHugh, 2012). We can observe from Fig 4(B) that at least 12 subjects are below 0.4 in the various scorer pairs again indicating the unreliable nature of scoring per subject among raters.

In particular the degree of disagreement was high for subjects that didn’t have a dominant alpha, thereby affecting the ability to rate the Hori scores as (1,2,3). For other subjects the degree of disagreement mainly arose due to the mislabelling of graphical elements. Examples of such typical cases of grapho elements are shown in Fig 4(C, D, E).

### 3.2. Automatic method

#### 3.2.1 External Validation: Spindle, K-complex detectors

The Spindle, K-complex detectors were validated externally using the DREAMS database along with other state of the art algorithms (Devuyst et al., 2011, 2010; Tsanas and Clifford, 2015) (detailed validation method in supplementary material). The validation results are shown in Fig 5. This validation ensured the element detectors perform on par with the state of the art methods. The parameters used in spindle, k-complex detectors (like spindle duration, k-complex amplitude etc.) were fixed with respect to the external databases and the same parameters were used in the validation of both Dataset #1, #2.

**Fig 5:**
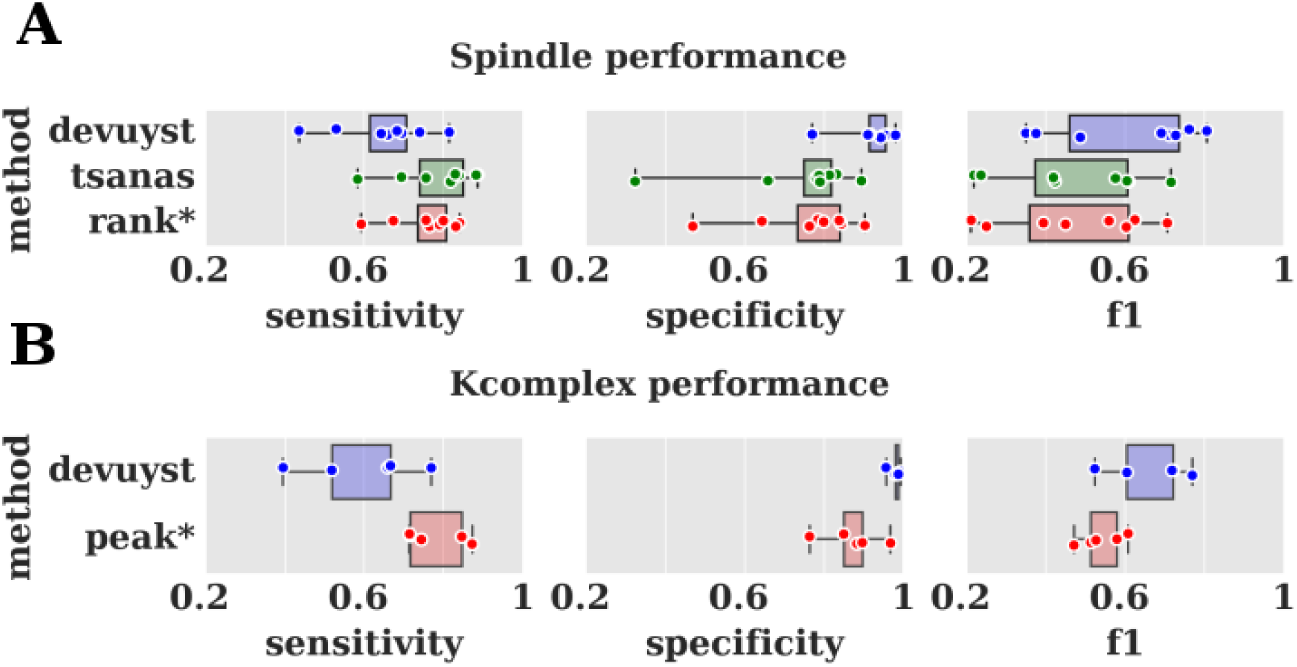
*Performance validation of grapho-element detectors with online database (DREAMS). The spindle detector was validated with state of the art algorithms from* (Devuyst et al., 2011; Tsanas and Clifford, 2015). The rank* algorithm developed in this paper performs comparable to the above mentioned algorithms. *The K-complex detector was validated with state of the art algorithms from* (Devuyst et al., 2010). The peak+ algorithm developed in this paper performs comparable to the above mentioned algorithms.

#### 3.2.2. Validation: Dataset#1

After the group consensus rule (sec 2.6.1) was applied on Dataset#1, the number of trials in the gold standard dataset in each class were: Alert:475, Drowsy(mild):1104, Drowsy(severe):281. Around 1306 trials (40%) did not have a consensus rating and hence were ignored from further analyses. This shows that about 40% of the overall trials didn’t have any consensus among the 3 different raters, further adding evidence to the disagreement among scorers mentioned in section 3.1.

Trials from all participants in Dataset#1 were first collated and then partitioned into 5 folds. The partition was made using stratified sampling such that the overall representation of subclasses remained similar in each fold. The training set further constituted of the first 4 folds and the test set consisted of the 5th fold. This procedure was repeated for 5 times as described in Fig 6(A). For each iteration the performance measures like sensitivity, specificity, f-1 scores were generated and the results are shown in Fig 7(A, B, C).

**Fig 6:**
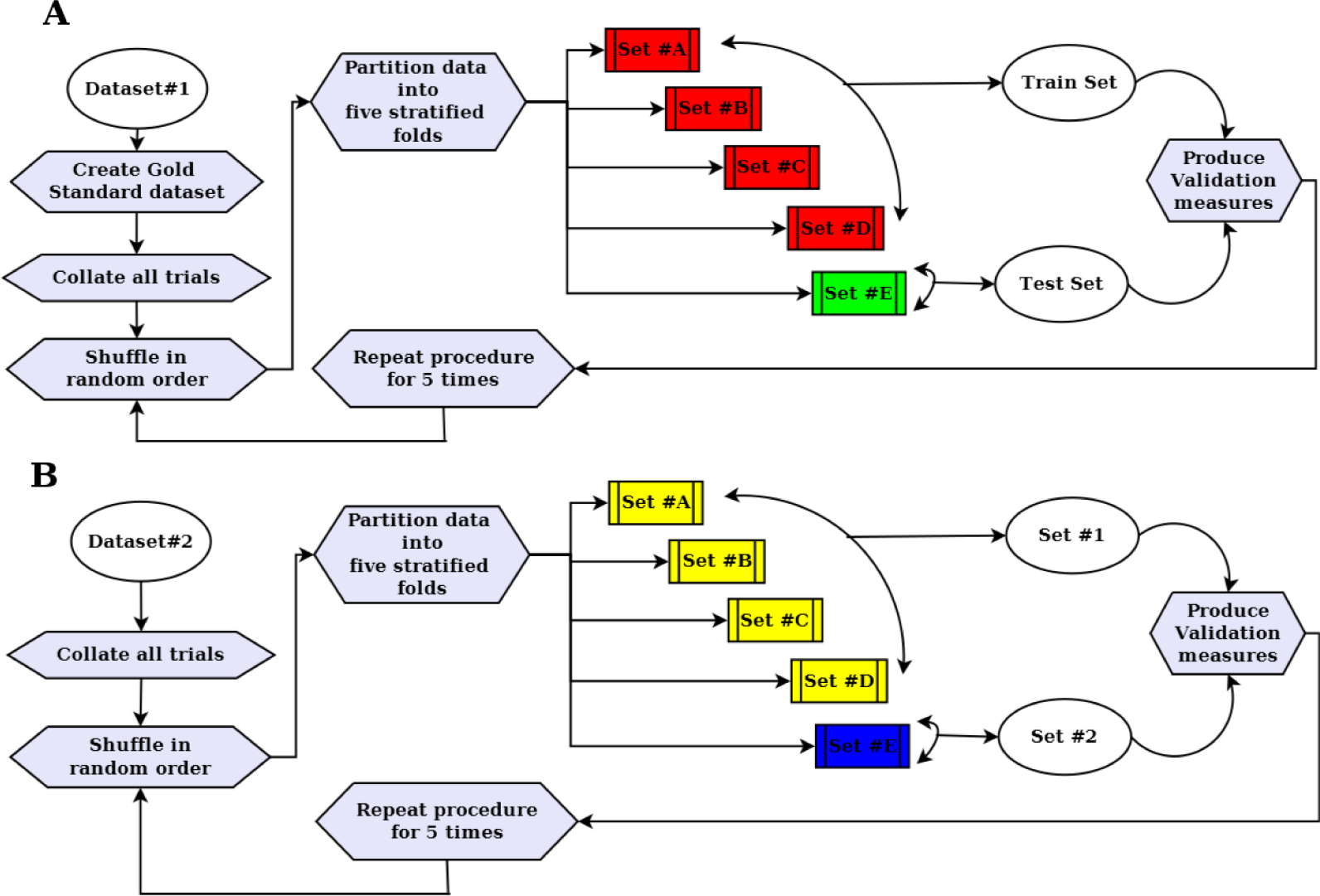
Curation of test and train datasets. (A) depicts creation of test and train dataset using Dataset #1 by five-fold stratified partition and this procedure is repeated for 5 times to produce validation measures. (B) depicts creation of Set #1, Set#2 using Dataset #2 by five-fold stratified partition and Set#1 is created by merging the first four sets and fifth set is constituted as Set #2 and this procedure is repeated for 5 times to produce validation measures.

**Fig 7:**
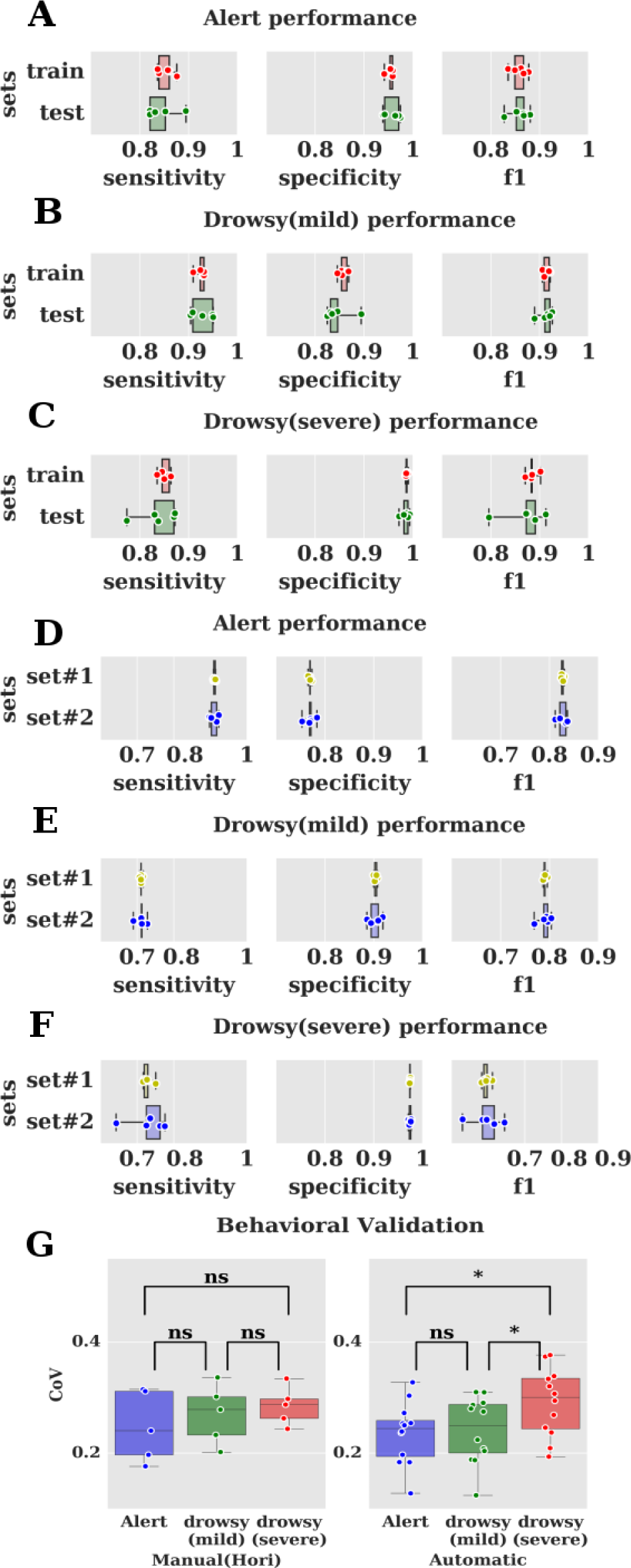
Validation measures of automatic algorithm. Validated with Dataset#1 using steps described in Fig 6(A). Results are depicted in the figure (A,B,C). The automatic algorithm was validated in an independent manner using Dataset#2 using steps described in Fig 6(B). Results are depicted in the figure (D,E,F). Validation with an independent measure (Coefficient of variation in reaction times) shows the algorithm reliably detecting differences (using repeated measures ANOVA) better than the manual scoring in figure G. ns: denotes p>0.05, * denotes p<0.01 (bonferroni corrected).

#### 3.2.3. Independent validation: Dataset#2

We decided to validate the algorithm (trained using dataset#1) on an independent dataset#2 to test its generalisability. This would mean that the hyper parameters (γ, C), support vectors trained using dataset#1 were directly applied on the dataset #2 without retraining. The number of trials in dataset#2 in each class were: Alert: 6049, Drowsy(mild): 7200, Drowsy(severe): 475. The dataset was divided into 5 folds using stratified sampling as before. The set#1 consisted of the first 4 folds and the set#2 consisted of the 5th fold. Thus set#1 contained atleast 4 times the number of trials in set#2 and hence similar in composition to the train and test sets in dataset #1 where train had at least 4 times the number of trials in test set. The same procedure was repeated 5 times as described in Fig 6(B). For each iteration the performance measures like sensitivity, specificity, and f-1 scores were generated and the results are shown in Fig 7(D, E, F).

The above mentioned methods in Dataset#2 tend to validate the automatic method against the human scorer. However, to claim that the automatic method out performs the human scorer in Dataset#2, we decided to further validate the same against an independent measure of drowsiness. Coefficient of variation (CoV) in reaction times has been used previously to measure drowsiness and is independent of both the observer and the algorithm’s pre-trial information (Bareham et al., 2014). We separated the trials among different classes of drowsiness using both the automatic and manual method. Further, CoVs were computed per participant for all classes (generated both by automatic and manual method) that contained at least 10 trials. Repeated measures ANOVAs on classes from automatic method yielded a main effect of drowsiness on CoV with F(2,22) = 9.25, p< 0.01. Post-hoc tests (Bonferroni corrected for multiple comparisons) yielded differences between mild and severe drowsiness (Cohen’s d: -0.95, p< 0.05), alert and severe drowsiness (Cohen’s d: -0.91, p< 0.05). However, the manual method failed to produce any main effect of drowsiness on CoV with F(2,8) = 1.2 with p> 0.05. These measures shown in Fig 7(G), clearly indicate the utility of the automatic method over manual scoring.

## 4. Discussions and Conclusions

In this paper, we have first described the pervasive problem of varying levels of alertness during cognitive experiments, particularly during eyes closed experiments. Such a scenario is further exacerbated in resting state EEG recordings. In many cases data from such experiments are used to compute measures like connectivity etc. that may further be contaminated by participants falling asleep (Tagliazucchi et al., 2012). This situation potentially contributes to wider problems faced by the scientific community such as the replication crisis.

In the past the problem of extreme relaxation and drowsiness has been sometimes ignored sometimes by cognitive scientists, who only take this confound into account by looking at reaction times and removing the sections where the participant was not responding or was too slow. Apart from visible changes in reaction times, there are changes in important processes like attention and perception as the participant drifts across varying levels of alertness (Goupil and Bekinschtein, 2012). Hence it is of paramount importance to control for varying levels of alertness. We have tried to solve this problem in an objective manner as follows. We first described the use of Hori scale that has been validated previously to detect the levels of alertness during the sleep onset process. However, the Hori scoring with 4 sec epochs is impractical to perform as it is highly subjective and time consuming (Ogilvie, 2001). In a typical experiment of about 600 trials well trained scorers take at least a day to score a single subject, and training new scorers takes atleast a month before they can be used for scoring. Using 3 independent raters on Dataset#1 we further quantified the inter-rater agreement using Krippendorff’s alpha and Cohen’s kappa metrics to show poor levels of agreement among the raters. This motivated us to develop an algorithmic solution that can be used to measure the level of alertness in a reliable manner.

Other attempts in the past to detect varying level of alertness using algorithms have suffered from several disadvantages. Firstly, such rule based algorithms (Olbrich et al., 2009) have validated their system using physiological measures like heart-rate variability etc. This further adds a layer of confound as measures of alertness need to be related again with physiological measures. Secondly, other algorithms (Crisler et al., 2008; Gudmundsson et al., 2005; Tagliazucchi et al., 2012) have been developed using traditional sleep stage based scoring. Such systems suffer from lack of resolution as they are validated with sleep scoring techniques that use 30 sec epochs. Thus they are unsuitable to match the micro dynamics in alertness observed during cognitive tasks. To our knowledge this is the first time an algorithmic solution has been attempted to measure the varying levels of alertness and simultaneously verified using a previously well validated system like Hori.

In the current work we have shown that predictor variance, coherence and grapho element detectors allow us to micro measure the level of alertness. We have constructed a classifier based on SVM and individual element detectors and have achieved sensitivity, specificity, f1-score of more than 0.8 in all subclasses (alert, drowsy (mild), drowsy (severe)) with respect to manual Hori scoring (gold standard from different raters). We have also validated our algorithm with a second independent dataset using different task conditions and recording electrode sites (using the same hyper parameters and support vectors trained using the first dataset). This produced a sensitivity, specificity, and f1-score of more than 0.7 in all subclasses. The main reason the performance reduces for drowsy (severe) subclass in dataset#2 is due to lack of gold standard comparison and fewer trials in this category. As the dataset#2 is scored only by one person it is prone to error (in a fashion similar to dataset#1 as depicted by varying levels of interrater agreement in Fig 4). This motivated us to show that our algorithm outperforms the manual scorer. Hence we employed a previously established independent behavioural measure of drowsiness using CoV in reaction times. We further showed that the automatic algorithm captures the variations in CoV better than the manual scorer in Fig 7(G). This stands testament to the generalisability of our method in detecting alertness levels across new datasets.

However, the use of Hori scale as validator has some disadvantages. Firstly, it is difficult to detect Hori stages (1-3) on participants who lack prominent alpha waves (Ogilvie, 2001). This would make these participants difficult to score manually, thereby explaining the lower sensitivity of the algorithm in the Drowsy (mild) subclass compared to the other classes. However, this is a problem for the human scorer, as the automatic algorithm is relatively immune to this problem, as it operates on relative variances across different bands rather than raw amplitude. Secondly, it has also been reported that the Hori stage (4) also doesn’t last long and hence is difficult to score (Ogilvie, 2001). Such samples would have had a high level of disagreement among scorers and hence would have been ignored while computing the gold standard dataset. Consequently, the difficult trials would not have been used for training the algorithm and hence it may not be able to detect any such trials in a new dataset. Thirdly, one of the main reasons for validating the algorithm with 3 subclasses is mainly due to lack of consensus in individual grapho elements. In order to truly validate the grapho elements we would need a dataset rich in those elements and also scorers who are able to consistently detect the grapho elements in a correct fashion.

The automatic algorithm devised here could be improved in several ways. Firstly, the current algorithm uses SVM with RBF kernels; other kernel choices like polynomial functions could be evaluated for making the optimal choice. Secondly, we performed only basic preprocessing of the pre trial data. However, it is well known that artifacts like eye movement, sweating, muscle artifacts can contribute to noise in the data. Hence the performance of the algorithm would improve if noise reduction measures are employed. However, we didn’t employ such measures as they are not standardized and we wanted to establish that the performance of the algorithm is robust under all conditions and hence performing specific pre-processing steps should not be an impediment for users of our method. Thirdly, we could also try to reduce the duration of epochs considered for labeling e.g. we can check the classification accuracies of signal durations of 1, 2, 3 secs etc. However, validating the same would be difficult as we also need to redo the human scoring with the corresponding reduced length of epochs. Fourthly, the automatic algorithm has been developed only for eyes closed condition. But many cognitive experiments have eyes open conditions and participants are also known to fall asleep under such active paradigms. The algorithm could be adapted for such paradigms; however detailed validation needs to be performed with other parallel measures of drowsiness like eye-tracking (as the Hori scale has not been validated for such purposes). Fifthly, the algorithm could further be refined to produce stages analogous to individual Hori stages. This would be helpful for researchers studying the sleep onset process in an objective manner as many complex non-linear changes in behaviour are known to occur in individual Hori stages (Noreika et al., 2017b). Finally, for quick paced experiments (short pre-trial periods), the parameters for detecting certain graphoelements (vertexes, k-complexes) are flexible to account for the shorter duration of the signal.

The applications of the algorithm include the following. Firstly, pre-trial data can be computed from task data (cognitive experiments) and the non-alert trials can be removed thus controlling for the effects of change in alertness levels. Secondly, we can detect and remove non-alert periods of data from resting state EEG experiments in a reliable manner. Thirdly, we can measure alertness as an independent variable and measure its effect on measures of interest. Fourthly, the method circumvents the subjective nature of the manual Hori scoring and thus enables to study the transition to sleep in an objective way. One of the most interesting aspects is the generalisability of the SVM classifier and other element detectors to the independent dataset#2, showing the high degree of transferability of this method, without having to retrain the classifier. Fifthly, when combined with online stimulus delivery techniques, the ability of our method to detect grapho elements (vertex, spindles, k-complexes) also allows us to investigate the effects of these elements on cognitive processes, for example by modulating the stimulus delivery according to the occurrence of these elements. Finally, sleep researchers can use this method for detecting N1 periods in the beginning of the night as well as awakenings and N1 periods during the full night period; further, they can also validate the detection of N2 periods by using the appearance of specific graphoelements (spindles, k-complexes).

All of the above mentioned facets make our method a powerful solution that can be used to micro measure varying alertness levels and thereby providing a valuable contribution to the study of both cognitive and resting state EEG experiments at large.

## Acknowledgements

This research was funded by Gates Cambridge Scholarship awarded to SRJ and Wellcome Trust Biomedical Research Fellowship WT093811MA awarded to TAB. We thank Louise Goupil for acquiring Dataset#1. We thank Valdas Noreika for providing Dataset#2 and many insightful comments on the manuscript, Carmen Soria for her work on a different manuscript on manual Hori scoring, We thank Srivas Chennu and Daniel Bor for their comments on a later version of the manuscript.

## Conflict of Interest

None

## Author Contributions

Conceptualization: SRJ, TAB

Data Curation: SRJ, TAB

Formal Analysis: SRJ, BJ, AEN, OVP, TAB

Funding Acquisition: TAB

Methodology: SRJ

Project Administration: TAB

Resources: TAB

Software: SRJ

Supervision: TAB

Validation: SRJ, CAB, BJ, AEN

Visualization: SRJ

Writing – original draft: SRJ

Writing – review & editing: SRJ, CAB, BJ, AEN, OVP, TAB

## 6. Supplementary methods

### 6.1. Vertex wave detectors

The two kinds of vertex waves depicted in Fig 2(B) are detected using the algorithm in Fig 8(A). As there was no online database available for vertex sharp waves it was not validated independently.

**Fig 8:**
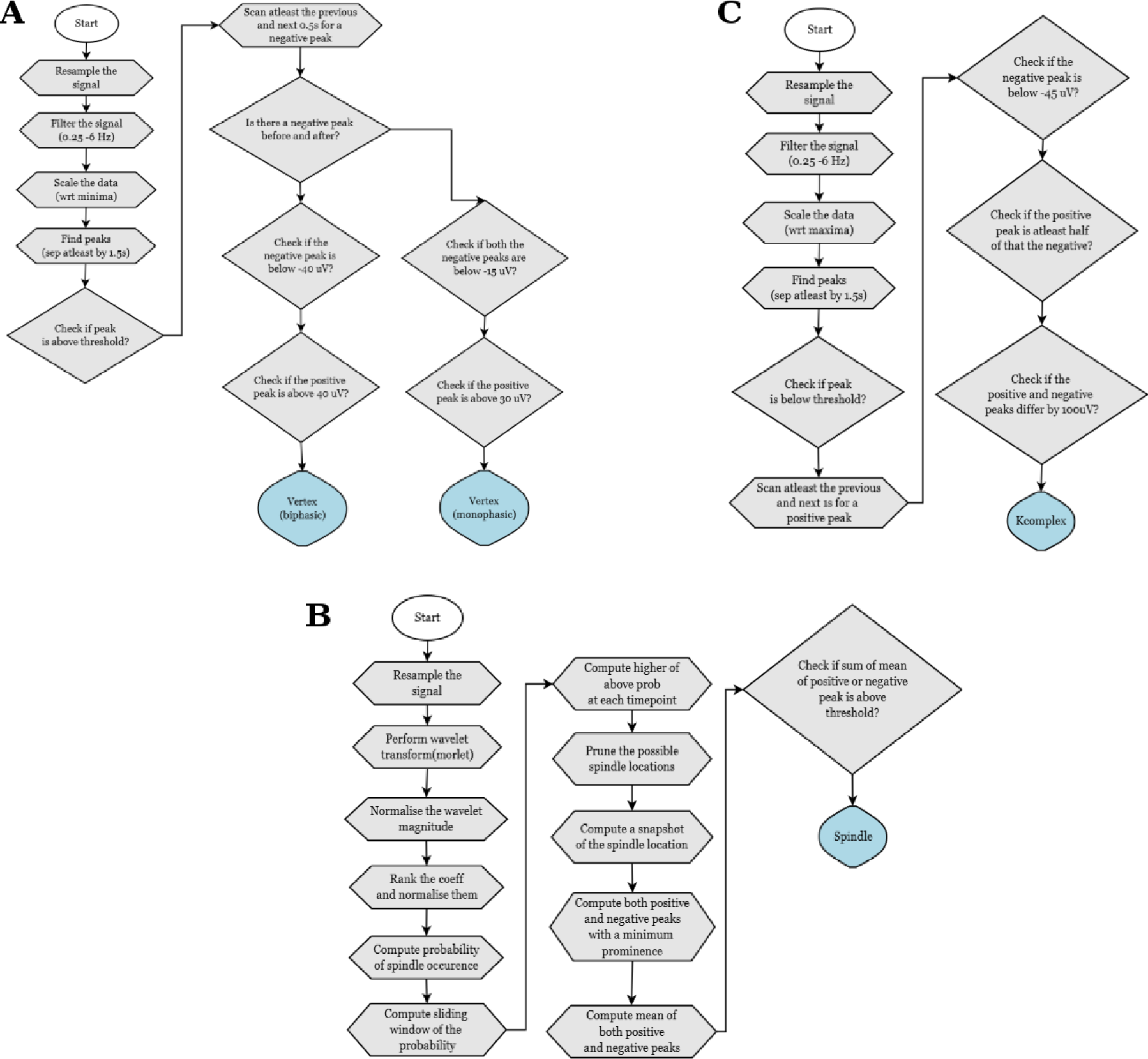
(A) Vertex wave detector algorithm. The preliminary step involves resampling, filtering and scaling of the signal to identify the peaks in the signal. Further the specific characteristics of the peaks are used to identify mono and biphasic vertex waves. (B) Spindle detector algorithm. The preliminary step involves resampling and using wavelet transform to identify the regions with high probability of occurrence of spindle waves. Further the specific characteristics of the waves are used to prune them. (C) K-complex detector algorithm. The preliminary step involves resampling, filtering and scaling of the signal to identify the peaks in the signal. Further the specific characteristics of the peaks are used to identify k-complex waves.

### 6.2. Spindle detectors

The spindles are detected using the algorithm in Fig 8(B). The algorithm was validated against an online database (DREAMS) (Devuyst et al., 2011) The data in the .edf format was first converted into EEGLAB format and was filtered from 0.5 - 20 Hz. The data was further resampled to 100 Hz and further epoched for each 4 sec. The gold standard dataset was created by merging the annotations from two experts for all the eight excerpts in the database. Our spindle detection algorithm was then validated against this gold standard along with state of the art methods that have already been validated against the same database (Devuyst et al., 2011; Tsanas and Clifford, 2015)

### 6.3. K-complex detectors

The Kcomplexes are detected using the algorithm in Fig 8(C). The approach developed here is similar (in terms of minima detection) to detectors developed elsewhere (Lajnef et al., 2015). The algorithm was validated against an online database (DREAMS) (Devuyst et al., 2010). The data in the .edf format was first converted into EEGLAB format and was filtered from 0.5 - 20 Hz. The data was further resampled to 100 Hz and further epoched for each 4 sec. The gold standard dataset was created by merging the annotations from two experts for the five excerpts in the database. Our kcomplex detection algorithm was then validated against this gold standard along with state of the art methods that have already been validated against the same database (Devuyst et al., 2010)

